# Divergent specializations for motion-driven representations in higher lateral and dorsal visual areas

**DOI:** 10.64898/2026.08.06.743321

**Authors:** Nastaran Darjani, Sophia Robert, Maryam Vaziri-Pashkam, Shahab Bakhtiari

## Abstract

The human visual system integrates both static and dynamic information to support form and shape perception, yet the computational principles underlying the integration of motion for object recognition remain unclear. Artificial neural networks (ANNs) offer a computational framework for developing and testing hypotheses about these principles: if ANNs trained on motion-related tasks develop representations that align with brain activity and support object categorization, this would suggest that the training objectives and architectural constraints of these networks may capture key aspects of motion processing in biological visual systems in general, and motion processing for object recognition, in particular. Here, we investigated this question using “object kinematograms”, stimuli in which object form is conveyed solely through motion cues. We measured neural responses of two higher regions of the lateral and the dorsal visual pathways, respectively, with strong sensitivity to dynamic cues from objects: lateral occipitotemporal cortex (LOT_bio_), and left supramarginal gyrus (SMG_lh_), as well as primary visual cortex (V1). We compared brain responses to representations extracted from two neural networks: SlowFast, a dual-pathway architecture trained on action recognition that processes slow- and fast-varying visual information with cross-pathway integration, and DorsalNet, a model of the primate dorsal visual pathway trained on embodied self-motion estimation. Representational similarity analysis revealed distinct representational profiles across brain areas, demonstrating functional specialization in motion-based form processing. LOT_bio_ was best characterized by the slow pathway of the SlowFast model, whereas SMG_lh_ showed strong similarity to both models. Critically, we found that representations aligned with brain activity also better supported behavioral function: the full SlowFast model, incorporating both slow and fast pathways, outperformed other models in few-shot categorization of object kinematograms and showed the highest similarity to human perceptual judgments. These findings demonstrate that with appropriate inductive biases, specifically, dual-pathway architectures for multi-scale motion processing and training objectives focused on dynamic visual tasks, ANNs can develop functionally useful representations of motion-defined forms that exhibit better alignment with the visual regions involved in processing dynamic visual signals.

## Introduction

Object recognition in natural environments requires integrating both static and dynamic visual information. Classically, these sources of information have been associated with two parallel cortical pathways: the ventral stream, specialized for object identity and form, and the dorsal stream, specialized for motion and spatial processing (1). In this traditional view, static features such as edges, textures, and shapes are primarily processed along the ventral pathway, whereas motion information is computed along the dorsal pathway and remains largely segregated from object representations. Following this principle, object recognition and motion processing largely developed as independent research problems. A large body of prior research has focused on static visual stimuli to understand object representations in the brain and in artificial neural networks (2–5). In contrast, computational studies of motion processing have primarily targeted dorsal regions and area MT, advancing our understanding of low-level motion extraction and self-motion estimation (6–8). More recently, however, this binary scheme has been extended by proposals of a third, lateral visual pathway. Projecting along the lateral occipitotemporal surface toward the superior temporal sulcus, this pathway has been argued to be specialized for the perception of dynamic and social stimuli, such as moving faces, bodies, and biological motion (9–11).

Beyond this proposed lateral specialization, the strict separation of form and motion across visual pathways does not appear to fully capture how the visual system operates (12, 13). Although relatively few studies have explicitly bridged object recognition and motion processing, emerging evidence suggests that motion information is also represented along the ventral pathway. For instance, dynamic stimuli have been used to probe object-selective cortex (11), and implied motion embedded in static images has been shown to modulate representations in both visual pathways (14). Furthermore, recent work demonstrates that object recognition processes extend beyond the ventral stream and involve contributions from dorsal cortical regions (10).

In line with this emerging view, Robert et al. (15) introduced “object kinematograms”, stimuli in which objects are defined purely by motion cues; and demonstrated that humans can reliably perceive and recognize motion-defined objects in the absence of static form information. In addition, these stimuli engaged higher-level occipito-temporal and parietal regions, including the lateral occipito-temporal cortex (LOT_bio_) and the left supramarginal gyrus (SMG_lh_), and revealed stronger decoding for dynamic compared to static stimuli in higher lateral areas. Together, these findings challenge the classical segregation of object recognition to ventral cortex and motion processing to dorsal cortex, indicating that both object- and motion-related representations are distributed across the visual pathways.

This convergence raises a central computational question: although motion information is present across pathways, do lateral and dorsal regions rely on similar computational principles, or do they implement fundamentally distinct computations? In other words, do their representational differences arise from separate goals, such as recognition versus self-motion estimation, or could they emerge from a common objective under different architectural or temporal constraints?

To address these questions, we adopted a model-based approach using artificial neural networks (16, 17). By examining which ANNs, with their specific training objectives and architectural constraints, best align with neural representations of motion-defined forms, we can generate testable hypotheses about the computations that support this capacity in the brain. Using object kinematograms (15) as stimuli, we probed the representational similarity of artificial neural networks (ANNs) with areas V1, LOT_bio_, and SMG_lh_. Given the observed role of these areas in representing object forms from motion cues, we turned to two types of neural networks with inductive biases for motion processing embedded in either their architectures or training objectives. The SlowFast network, trained to perform action recognition from video data, has parallel pathways, each operating at different temporal sampling rates (18). The presence of the two pathways allows the extraction of both fast- and slow-varying visual features, enabling us to measure the contribution of fast- and slow-varying information in representational similarity with our brain areas of interest. In addition, we compared brain responses to the DorsalNet model (6), which was directly trained to estimate self-motion parameters from egocentric video inputs. Together, these two models allow us to examine whether human cortical regions are best described by categorical motion-based object processing, low-level motion feature extraction, or a combination of both.

By comparing representational dissimilarity matrices (RDMs) from fMRI with those derived from SlowFast and DorsalNet, and by manipulating pathway interactions, we sought to answer four key questions: (1) how temporal scales of motion are differentially represented across cortical regions, (2) whether higher-order visual areas are better captured by models that integrate slow and fast motion cues, (3) whether lateral and dorsal regions implement motion-based object representations for different functional objectives, and (4) to what extent computational models account for the behavioral relevance of motion-based object recognition. Our analyses revealed distinct patterns of model-brain correspondence across higher visual areas, suggesting that different aspects of motion processing may contribute to representations in these regions. In addition, the model with representational geometry most similar to that from human similarity ratings showed the best few-shot categorization learning on object kinematogram stimuli, suggesting the sample efficiency of the representations underlying human perception in learning motion-defined object categories.

## Results

In this study, we investigated how the brain encodes motion-defined form and shape by linking neural representations across visual areas with computational models optimized for dynamic vision. To this end, we used object kinematograms, a type of stimulus in which object form is defined solely by motion (15). fMRI data were collected from V1, LOT_bio_, and the left SMG while participants viewed these stimuli (Fig. 1A). For each region, responses were summarized as category-level representational dissimilarity matrices (RDMs; 6*×* 6, computed after averaging across stimulus exemplars within each of the six object categories; see Methods). These brain RDMs were then compared with RDMs derived in the same way from two neural network architectures: SlowFast, a dual-pathway model integrating slow and fast temporal streams (18), and DorsalNet, a model designed to capture self-motion features within the dorsal pathway (6) (Fig. 1B and C).

**Fig 1.**
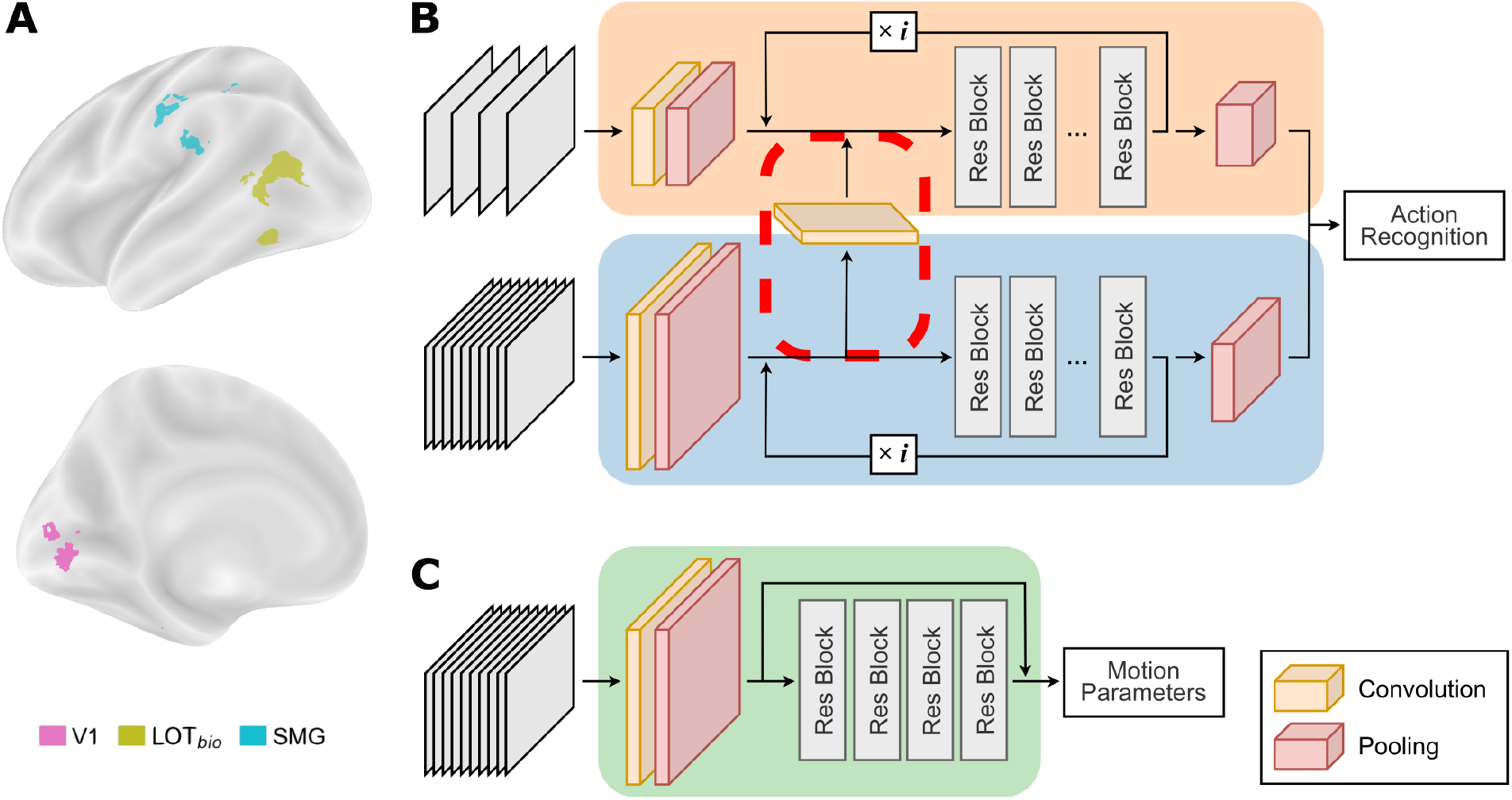
A.Regions of interest (ROIs) for one example subject. ROIs are shown in distinct colors, as indicated in the legend. B. Schematic of the SlowFast network. The orange pathway represents the slow stream, and the blue pathway represents the fast stream. The red dotted line highlights the cross-connection between the two pathways. In the schematic, i denotes the iteration of the module and was set to 4 in the model, and the number of residual blocks in each iteration indicated by Nres(i) = [3, 4, 6, 3]i. C. Schematic of DorsalNet, a motion-sensitive network predicting self-motion parameters by mimicking the computational properties of the primate dorsal visual stream.

### Representational similarity between brain regions and models

We first examined the correspondence between neural representations in V1, LOT_bio_, and the SMG_lh_ with those derived from the SlowFast and DorsalNet models (Fig. 2). Consistent with its role in low-level visual encoding, V1 showed little to no correspondence with any model layer, and the near zero noise ceiling value limited the interpretability of further analyses. In contrast, higher-level regions showed distinct and robust patterns of similarity. LOT_bio_ aligned most strongly with the slow pathway of SlowFast (S_wx_: the slow pathway with cross-connection from the fast pathway), particularly in its deeper layers (peak *τ* = 0.25; cluster-based permutation test, *N* = 1000, *p* < 0.05, corrected across layers). Note that S_wx_ integrates information from the Fast pathway via cross-pathway connections (Fig. 1B), unlike the Fast pathway itself, which operates in isolation. The SMG_lh_ also exhibited significant alignment with S_wx_ (peak *τ* = 0.23; cluster-based permutation test, *N* = 1000, *p* < 0.05, corrected across layers).

**Fig 2.**
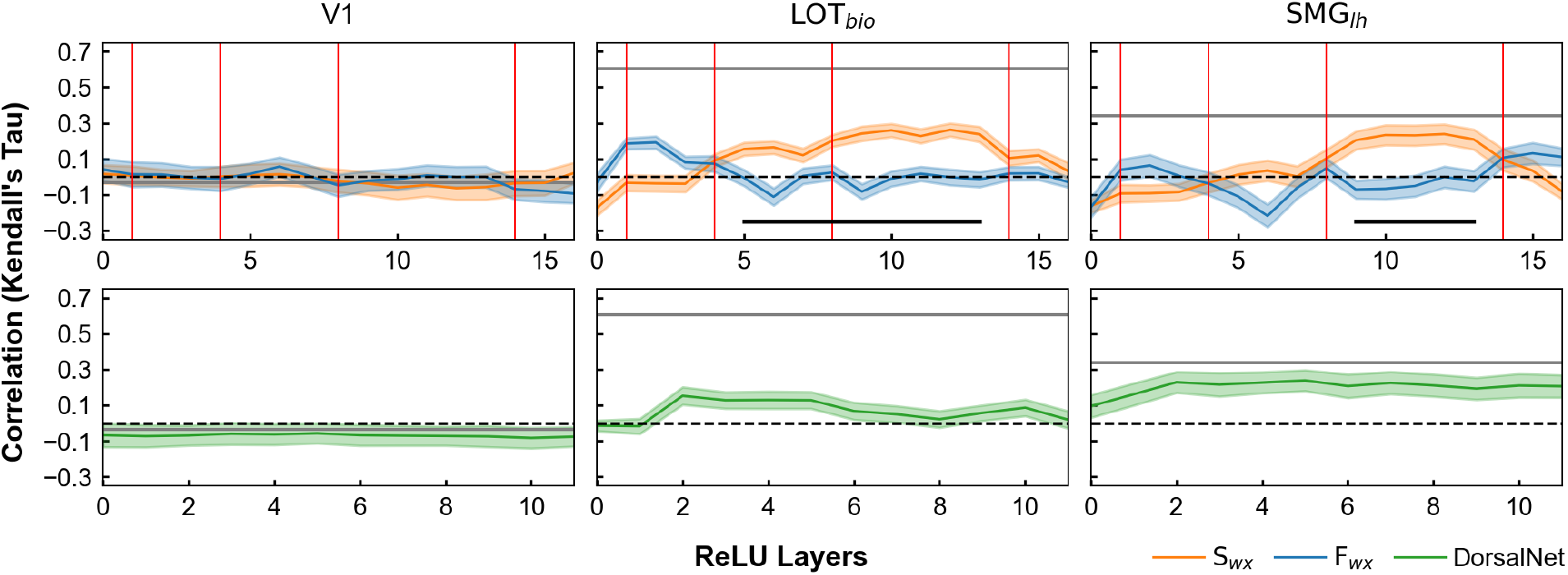
Representational similarity analysis between fMRI regions and model layers for the SlowFast (top) and DorsalNet (bottom) architectures. S_wx_ and F_wx_ denote the slow and fast pathways of the SlowFast model. The gray line indicates the noise ceiling (mean subject reliability). Red vertical lines mark position of cross-pathway fusion layers, and black lines indicate layers with significant difference between models (cluster permutation test, p < 0.05). Shaded areas indicate the SEM across subjects.

When compared to DorsalNet, a different profile emerged (Fig. 2, bottom). In SMG_lh_, correlations with DorsalNet layers were strikingly high, approaching the noise ceiling from the earliest stages through the final layers (peak *τ* = 0.22; cluster-based permutation test, *N* = 1000, *p* < 0.05, corrected across layers), suggesting that this region is well described by models optimized for self-motion and embodied visual prediction rather than categorical action recognition. Since DorsalNet was designed to model primate dorsal visual pathways through self-motion estimation during navigation, this pattern suggests that SMG_lh_ representations are more similar to those learned for embodied visual processing and self-motion inference than those in LOT_bio_.

Since both S_wx_ and DorsalNet show high correspondence with SMG_lh_, we next asked whether they capture overlapping or distinct aspects of brain representations. To address this, we performed partial correlation analyses on the highest-correlated layers of each network. In SMG_lh_, correlations with S_wx_ (*τ* = 0.23, *τ*_partial_ = 0.14 *±* 0.04) and DorsalNet (*τ* = 0.22, *τ*_partial_ = 0.13 *±* 0.05) were largely shared, indicating substantial overlap in the motion-related information captured by the two models. At the same time, each network retained unique variance, suggesting that although both align well with SMG_lh_, they emphasize partially distinct representational features. In contrast, LOT_bio_ (*τ* = 0.25, *τ*_partial_ = 0.21 *±* 0.03) retained substantial unique variance with S_wx_ even after controlling for DorsalNet, indicating that the SlowFast model captures aspects of LOT_bio_ representations that are not explained by DorsalNet (Fig. 3). Together, these findings suggest that the two higher-order regions exhibit different patterns of correspondence with the computational models. Whereas LOT_bio_ is more selectively aligned with the temporally integrated representations of the SlowFast model, SMG_lh_ shows substantial correspondence with both SlowFast and DorsalNet, indicating that each model captures complementary aspects of the representational geometry in this region.

**Fig 3.**
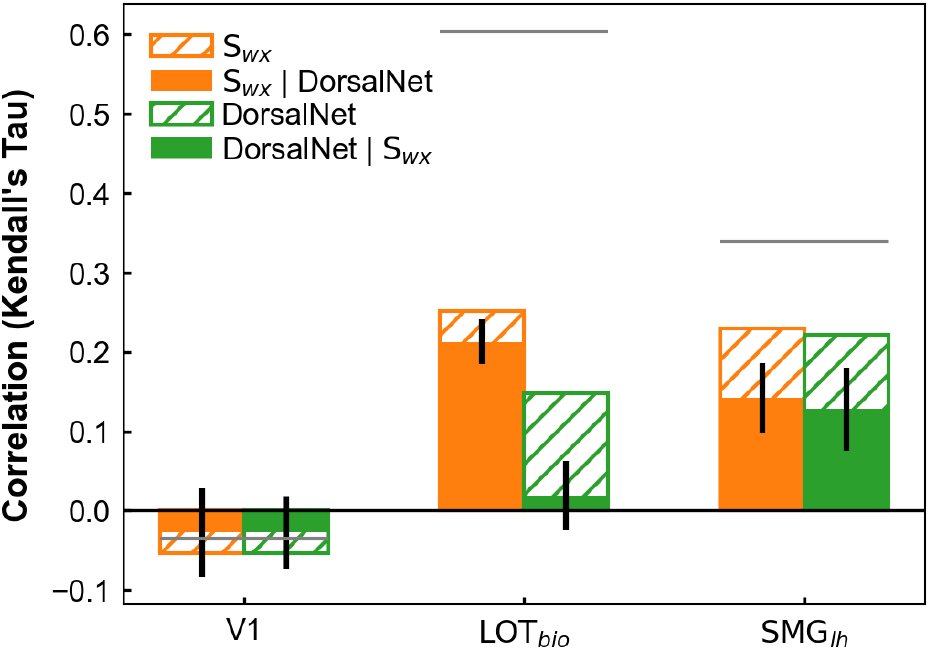
Zero-order (hatched) and partial (filled) correlations of the most predictive layers from S_wx_ and DorsalNet with fMRI regions. Partial correlations control for variance explained by the other model. The gray line indicates the noise ceiling, and error bars represent SEM across subjects in partial correlations.

### Role of multi-scale temporal integration and cross-pathway connections

The SlowFast architecture comprises two pathways operating at different temporal resolutions: a slow pathway that captures coarse, slowly varying motion cues, and a fast pathway that processes fine, rapid temporal changes. Crucially, cross-pathway connections allow information from the fast pathway to feed into the slow pathway, enabling temporal integration across timescales (Fig. 1B). Examining the role of these connections allows us to evaluate whether integrating information across multiple temporal scales improves the correspondence between model and brain representations.

To probe the role of temporal integration, we compared the intact SlowFast slow pathway with (S_wx_) and without (S_nox_) cross-pathway connections (Fig. 4 shows the correlation of highest-correlated layers of each network). In LOT_bio_, removing cross-connections had little impact, suggesting that the fast-varying information input to the slow pathway through the cross-connection does not play a significant role in the high similarity of LOT_bio_ with S_wx_. In SMG_lh_, however, similarity drastically decreased when cross-connections were absent, confirming that the similarity of this region with the network depends on the integration of fast-to-slow information.

**Fig 4.**
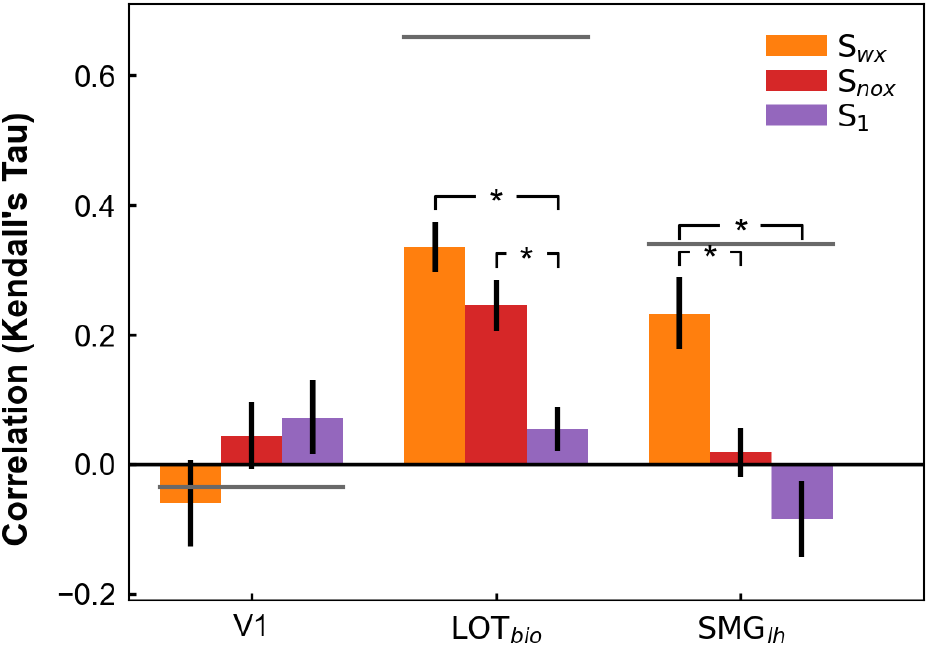
Representational similarity of SlowFast pathway variants with brain regions at layer 10. S_wx_ represents the intact slow pathway, S_nox_ the slow pathway without cross-connections, and S_1_ an independently trained slow-only model. The gray line indicates the noise ceiling. Asterisks denote significant differences between models at layer 10 (i.e., whether this layer falls within a significant cluster, cluster permutation test, p < 0.05). Error bars show SEM across subjects.

Even after ablating the cross-pathway connection, the slow pathway still had traces of sensitivity to fast-varying information due to the presence of the cross-pathway connection during training, as we only remove cross-connections after training the network on its action recognition task. Therefore, we investigated whether exposure to the fast pathway’s motion during training changed the representational geometry of the slow pathway. To this end, we compared S_nox_ with a pure slow-only model trained independently (S_1_). LOT_bio_ showed higher similarity to S_nox_ than to S_1_ (*τ*_Snox_ = 0.24 *±* 0.04, *τ*_S1_ = 0.05 *±* 0.03 for LOT_bio_; cluster-based permutation test, *N* = 1000, *p* < 0.05, corrected across layers). This indicates that even if fast cues were not directly required at inference for LOT-like representation, the presence of fast-varying information during training was essential for shaping representations similar to LOT_bio_. In SMG_lh_, both the slow pathway without cross-pathway connections (S_nox_) and the independently trained slow-only model (S_1_) showed significantly lower similarity to SMG_lh_ compared with the intact model (S_wx_) (*τ*_Swx_ = 0.24 *±* 0.06, *τ*_Snox_ = 0.01 *±* 0.05, *τ*_S1_ = -0.09 *±* 0.06; cluster-based permutation test, *N* = 1000, *p* < 0.05, corrected across layers). This indicates that SMG-like representations depend on the integration of slow- and fast-varying information both while the representation is being learned and at inference, rather than on fast-varying information alone.

### Representational structure of brain–model relationships

We ran a multidimensional scaling (MDS) analysis to obtain an overall view of representational relationships of brain regions and models that we included in our study. For each participant, we first computed the representational similarity between all pairs of brain regions (V1, LOT_bio_, and SMG_lh_) and computational models (S_wx_, F_wx_, S_nox_, S_1_ and DorsalNet) based on their RDMs. These similarity values were then projected into joint reduced 2D representation across participants using MDS.

The resulting two-dimensional visualization (Fig. 5) shows the positions of each human subject’s brain region (small colored circles), the average regional cluster positions across subjects (large colored circles), as well as each ANN model included in our analysis (empty large circles). The analysis revealed that S_wx_ clustered closely with LOT_bio_ and SMG_lh_, suggesting it as the most brain-like component of the SlowFast architecture. DorsalNet was positioned closer to SMG_lh_ than to the other regions, consistent with the RSA results in the previous section. In contrast, V1 exhibited a highly dispersed cluster, reflecting the variability of its representational geometry. This pattern likely arises from V1’s inability to integrate the global motion patterns that define objects in the kinematograms (given the small receptive field of V1 neurons). As a result, V1 showed both a low noise ceiling in the earlier analyses and a dispersed representation in the current MDS space.

**Fig 5.**
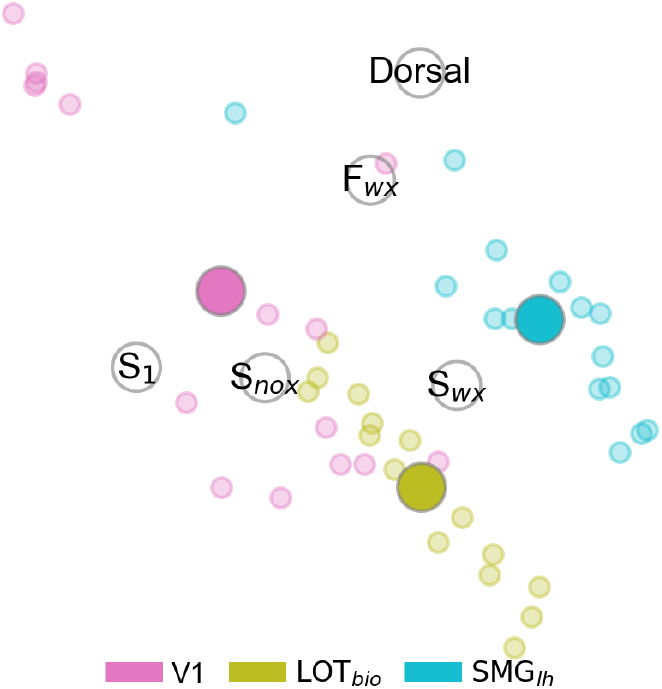
Two-dimensional multidimensional scaling (MDS) visualizing the representational similarities between brain regions and models. Small points represent individual participants; large filled circles indicate group-average positions.

The MDS embedding also revealed a pattern within the SlowFast family. The independently trained slow-only network (S_1_) was positioned farthest from the brain regions, but adding higher-frequency motion information during training (S_nox_) shifted the model closer to neural data. Eventually, the integration of fast and slow pathways (S_wx_) further improved alignment, particularly for SMG_lh_, while the fast pathway alone (F_wx_) occupied a position closer to DorsalNet, capturing complementary information about motion features. This progression illustrates how architectural integration and motion training progressively align the models’ representations with those of the visual regions we examined.

### Behavioral relevance of motion-based object representations

The results so far demonstrate how the integration of slow- and fast-varying information is crucial for reproducing brain-like representational geometry in response to motion-only features (object kinematograms). Finally, we assessed whether the networks could perform object categorization based on object kinematograms, as humans can do. To this end, we added a classification head to each ANN model to categorize the six stimulus classes used in the fMRI experiment (human, mammal, reptile, tool, pendulum/swing, and ball) while freezing all other layers. We trained each classification head of the models by augmenting the training set by varying random seeds for object kinematogram generation. Using this approach, the model only observed 10 object instances per category during fine-tuning. Random seed initializations did not increase the number of instances but only added noise diversity to the stimuli. Testing was performed on the original stimuli, to match those used in the fMRI experiment. Both S_1_ and S_nox_ performed poorly, failing to reliably classify the six object categories (acc_S1_ = 15.56 *±* 1.11%, acc_Snox_ = 17.78 *±* 0.68%). DorsalNet achieved higher than chance accuracy similar to the fast pathway in the SlowFast model (acc_dorsal_ = 26.11 *±* 2.42%, acc_Fwx_ = 25 *±* 1.52%). Higher performance was achieved by the Slow pathway with cross-connections (acc = 36.67 *±* 2.04%). In addition, combining and entangling both slow and fast information with cross-connection significantly outperformed all other models (acc = 47.22 *±* 3.62%), including both S_wx_ alone and F_wx_ alone, each pair compared directly using a permutation test on the accuracy difference (all pairwise comparisons *p* < 0.05; Fig. 6).

**Fig 6.**
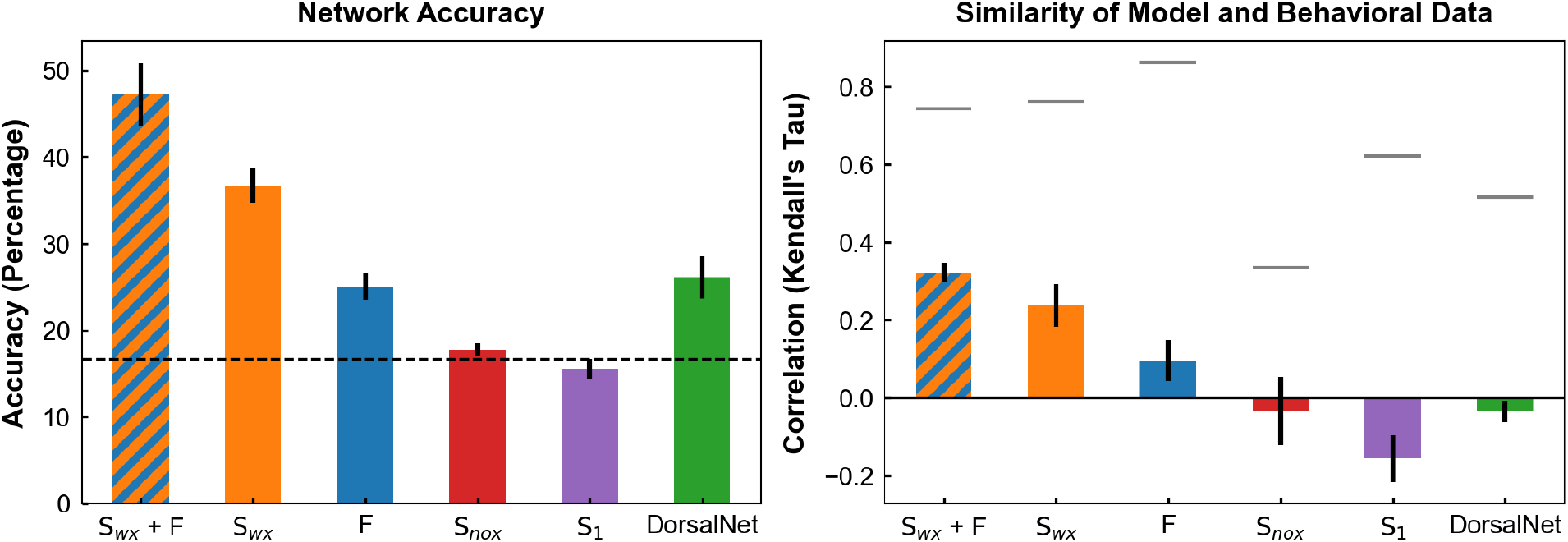
Classification accuracy of each model on the object kinematogram task. Dotted line indicates chance (left panel). Correspondence between model confusion matrices and behavioral RDMs. The gray line marks the noise ceiling, and error bars show SEM across folds (right panel).

We next compared the confusion matrices from model predictions with the behavioral representational dissimilarity matrix reported by Robert et al. (15). Two ablations of the SlowFast model, which alter information flow from the fast pathway, did not match behavioral data (*τ*_Snox_= -0.04 *±* 0.09, *τ*_S1_ = -0.16 *±* 0.06). Surprisingly, DorsalNet showed little to no similarity to human behavior (*τ* = -0.04 *±* 0.03), despite its strong alignment with SMG_lh_ neural responses. By contrast, the slow and fast pathways of the SlowFast model exhibited positive correlations with behavior, with the slow pathway showing significantly stronger similarity than the fast pathway (*τ*_Swx_= 0.24 *±* 0.06 vs. *τ*_Fwx_ = 0.096 *±* 0.05; permutation test on the correlation difference, *p* < 0.05). In addition, the combined S_wx_*+ F* pathway (the intact output of SlowFast network) achieved the highest similarity to human behavior, significantly outperforming all other models (*τ* = 0.32 *±* 0.03), including both S_wx_ alone and F_wx_ alone (each pair compared directly with a permutation test on the correlation difference, all *p* < 0.05). Together, these results highlight two key findings: first, the behavioral task used here is better captured by networks optimized for object and action recognition than by a network optimized for self-motion estimation and embodied visual processing, as evidenced by the poor behavioral alignment of DorsalNet; and second, within such networks, integrating slow- and fast-varying motion information yields substantially closer similarity to human behavior than either temporal scale in isolation.

## Discussion

In this study, we investigated motion-defined object representations in two higher-order regions that preferentially respond to dynamic stimuli: LOT_bio_, a region associated with the lateral visual pathway, and SMG_lh_, a parietal region of the dorsal visual pathway. We examined how these representations relate to artificial neural networks optimized for different computational goals. Specifically, we compared neural representations to two artificial neural network models: a two-pathway architecture optimized for action recognition that separates slowly and rapidly varying motion dynamics (18), and DorsalNet, a model trained to estimate self-motion from visual input, thereby learning representations that support navigation and embodied visual understanding (6). Through this representational comparison, we asked whether the training objectives and architectural inductive biases of these models provide insight into the representational properties of these higher-order ventral and dorsal visual areas. Focusing on V1 as a baseline, and on LOT_bio_ and left SMG as representative motion-sensitive ventral and dorsal regions, respectively, our results extend recent findings that motion-defined object perception engages a distributed cortical network (15). Here, we show that the two higher-order regions, despite their shared sensitivity to motion, differ in the nature of their motion representations and in how these representations may support distinct downstream perceptual and behavioral functions.

Our results demonstrated that, consistent with its role in early visual processing, V1 showed no reliable similarity to any of the tested models, reflecting its limited capacity to integrate global motion structure in object kinematograms. In contrast, higher-order regions exhibited systematic and dissociable alignment patterns. The lateral region, LOT_bio_, was best captured by the slow pathway of the SlowFast model, particularly when that pathway was trained with access to fast-varying motion information (i.e., fast and slow pathways trained together with cross-pathway connections). In contrast, SMG_lh_ showed strong correspondence with both the SlowFast model and DorsalNet, with each model explaining a unique component of the representational structure in this region.

Second-order representational analyses, conducted via estimating pairwise similarities between the representational dissimilarity matrices (RDMs) of brain regions and computational models, revealed that increasing exposure to motion statistics and cross-pathway integration progressively shifted model representations toward those observed in the human cortex (Fig. 5). In a low-dimensional embedding of representational geometries, the SlowFast variant lacking access to fast temporally sampled motion information (S1) was positioned farthest from neural data, whereas incorporating faster motion statistics to the model moved representations closer to cortical organization. Consistent with the representational similarity analyses, DorsalNet and the fast pathway of the SlowFast model were positioned closer to SMG_lh_ than to LOT_bio_ in the MDS space. In contrast, the fully integrated SlowFast model (S_wx_) occupied a position between LOT_bio_ and SMG_lh_ and showed closer correspondence with both regions than the other SlowFast variants.

Importantly, this same model also best predicted human behavioral performance (Fig. 6), indicating that successful object perception depends not simply on motion sensitivity, but on how motion cues are integrated over time to support object-relevant representations. This convergence across neural and behavioral data highlights the functional importance of multi-scale integration of motion processing.

### Effect of Temporal Resolution on Motion-Based Object Representations

The SlowFast architecture provided a principled computational framework for examining how different temporal sampling regimes contribute to perceiving motion-defined objects. Both different pathways in this architecture process dynamic information, but in different temporal resolutions. Moreover, by incorporating cross-connections between the two pathways, the model can capture how the integration of signals at different temporal resolutions impacts alignment with the brain.

Our results suggest that motion information contributes to representations in both ventral and dorsal visual areas, but the reliance on temporal scales differs. In the dorsal region (left SMG), the sharp reduction in alignment with SlowFast following ablation of fast temporal sampling (Fig. 4) suggests that SMG_lh_ depends critically on multiscale temporal information and integrates complementary motion representations across slow and fast timescales.

In the ventral stream (LOT_bio_), slow-varying motion supports the stability required for object recognition. However, we found that LOT_bio_ representations align more closely with a SlowFast architecture than a Slow-only model. This discrepancy suggests that representations learned with exposure to fast-varying motion during training exhibit greater correspondence with LOT_bio_ than representations learned without such exposure. Consequently, the representational geometry of ventral regions may be an indirect byproduct of fast temporal statistics encountered during learning.

A related theoretical perspective is provided by Slow Feature Analysis (19), which proposes that slowly varying features extracted from dynamic input provide a stable basis for invariant visual representations, such as object identity. While this framework aligns with our observation that slow-varying information contributes to stable representations, our results point to a more nuanced picture. Specifically, we find that to obtain dorsal-like representations, fast-varying motion information is critical. Also, to obtain ventral-like representations, it is necessary to have fast-varying motion information present during learning. These findings suggest that visual representations are shaped not only by slow temporal components, but by the interaction between multiple temporal scales, providing a complementary extension to their SFA framework.

### Effect of Computational Objective on Motion-Based Object Representations

Although motion information is ubiquitous across cortical pathways, how it is represented is fundamentally shaped by the computational goals of each region. In LOT_bio_, representations were better predicted by SlowFast than by DorsalNet. This suggests that ventral/lateral motion re-presentations are not shaped by self-motion estimation – the objective on which DorsalNet was trained – but instead support categorical object and action representations, consistent with the established role of the ventral/lateral visual pathway. By contrast, SMG_lh_ exhibited a hybrid profile, with both DorsalNet and SlowFast capturing significant variance. Together, these patterns point to a divergent specialization: lateral motion representations are tied selectively to recognition-oriented objectives, whereas dorsal representations reflect both recognition- and self-motion embodied computations. It is worth noting, however, that LOT_bio_ and SMG_lh_ were not defined using identical criteria: LOT_bio_ was functionally localized for selectivity to biological motion, whereas SMG_lh_ was defined anatomically from all visually responsive voxels in the region. This difference in ROI definition could itself contribute to the distinct representational profiles we observed, and future work using functionally matched localization criteria across regions would help clarify whether the dissociations reported here reflect genuine differences in computational objective rather than differences in the specificity of the underlying voxel populations.

While we have characterized LOT_bio_ and SMG_lh_ through the lens of different computational goals, these specializations might alternatively emerge from a single objective function, such as self-supervised prediction. Previous work in mice (7) and humans (12) suggests that ANNs optimized for prediction develop a variety of visual specializations that mirror the dorsal-ventral split. Consequently, the distinct motion representations we found in LOT_bio_ and SMG_lh_ may not reflect optimization for isolated tasks, but rather the internal heterogeneity that arises when a single system is trained to predict future visual states. Distinguishing these two accounts – divergent objectives versus a shared predictive objective with emergent specialization – is an important direction for future work.

### Behavioral Validation of Motion-Based Representations

If computational objectives shape the representational geometry observed in ventral and dorsal cortex, a critical test is whether these same objectives determine how well models account for human perceptual behavior. The behavioral analyses provide such a test by evaluating whether motion-based representations learned under different optimization goals support object categorization and reproduce the similarity structure observed in human judgments. Across models, the results revealed a graded pattern that closely mirrored both architectural differences and training objectives.

First, the behavioral task examined here, object categorization and similarity judgments based on motion-defined objects, was better captured by the SlowFast model than by DorsalNet. Although DorsalNet aligned strongly with SMG_lh_ neural responses, it failed to capture the structure of human similarity judgments and showed only modest above-chance categorization performance. This dissociation likely reflects distinct optimization objectives of the models rather than the presence or absence of motion representations. Both SlowFast and DorsalNet learn motion-based features, but SlowFast is optimized for action and object-related recognition tasks, whereas DorsalNet is optimized for self-motion estimation and embodied visual processing. Because the behavioral measures used here emphasize object-level perceptual judgments, they may more closely reflect the representational properties of lateral occipito-temporal cortex than those of SMG_lh_. This interpretation is consistent with evidence that behavioral similarity judgments are more strongly associated with representations in lateral occipito-temporal action networks, whereas parietal action representations are more strongly related to body-part movement information, suggesting that different motion-processing systems support distinct representational goals (20).

A similar limitation was observed for SlowFast variants lacking effective temporal integration. Slow-only representations without proper exposure to richer temporal statistics (S_nox_, S_1_) performed at chance level and showed zero or negative correlation with behavioral similarity structure. These findings suggest that temporally slow representations alone cannot support motion-defined object recognition.

In contrast, models optimized for action recognition demonstrated progressively stronger behavioral correspondence when equipped with appropriate temporal resolution and cross-pathway integration. By introducing fast-varying information to networks, the results for both categorization accuracy and human behavioral similarity, significantly improved. The SlowFast architecture, particularly its slow pathway, supported higher few-shot categorization accuracy, consistent with the idea that slowly varying, temporally integrated motion cues provide stable information for object-level judgments. The strongest performance was achieved by the fully integrated SlowFast model combining S_wx_ with the Fast pathway, demonstrating that interaction between slow and fast dynamics yields the most behaviorally effective representations.

Taken together, these results suggest that temporal integration plays an important role in shaping motion-based object representations that align with both neural and behavioral organization. Models incorporating interactions between slow and fast temporal pathways showed stronger correspondence with human behavioral similarity judgments and higher alignment with higher-level visual regions, whereas models trained primarily for self-motion estimation or lacking cross-pathway integration showed weaker behavioral correspondence. Although the SlowFast architecture was originally optimized for action recognition rather than explicit object categorization, its integration of information across multiple temporal scales appears to support representations that are more consistent with human perceptual organization.

### Rethinking the static–motion dichotomy in visual pathways

Despite recent advances, most contemporary ANN models of vision remain limited in their ability to capture the richness of human visual representations, particularly with respect to the integration of motion and form. Models of core object recognition have largely focused on static images (2–5). While highly successful for explaining ventral-stream responses to static stimuli, these models typically disregard motion information and therefore provide an incomplete account of visual processing in dynamic environments.

In parallel, a smaller body of work has focused on motion processing, characterizing how preferences for simple to increasingly complex motion features emerge across multiple stages of processing. Although these models have been valuable for explaining motion-selective responses in dorsal regions, they are generally optimized for self-motion estimation and pay little attention to object identity or form (6–8). As a result, both modeling traditions: static object recognition on the one hand and motion processing on the other, have advanced largely in isolation, each capturing only a subset of the representations present in the human visual system.

This study focuses on motion information processing over two motion-sensitive regions, one ventral and one dorsal, and thereby aims to bridge this isolation. By using motion-isolated stimuli, we show that in the absence of appearance cues, the geometrical similarity of cortical responses is primarily driven by differences in motion representations. Comparing two networks with distinct architectures and optimization objectives further clarifies how motion is utilized across pathways. Overall our results challenge the classic dichotomy that assigns static object recognition to the ventral stream and motion processing exclusively to the dorsal stream (1, 21). Instead, motion information and object recognition are possibly processed throughout the visual hierarchy in an entangled manner (7, 10, 12, 15). The critical distinction lies not in whether motion is processed, but in how it is computed and represented to support perceptual goals. Integration across temporal scales, rather than a strict dorsal–ventral division, appears central to dynamic vision.

## Methods

### Participants and fMRI dataset

We used the publicly available fMRI dataset from Robert et al. (15), in which participants viewed motion-defined object stimuli during a passive viewing task. The dataset included responses from 15 participants across multiple visual and parietal regions of interest (ROIs). Based on their findings, we focused on three ROIs: primary visual cortex (V1), lateral occipito-temporal cortex selective for biological motion (LOT_bio_), and the left supramarginal gyrus (SMG_lh_). The regions were defined following the atlas- and localizer-based approach reported by Robert et al. (15). V1 was included as a baseline region, while LOT_bio_ and SMG_lh_ were selected to represent ventral and dorsal pathway regions, respectively, both of which show greater sensitivity to dynamic than to static stimuli (15). Notably, the two regions were not defined using identical criteria: LOT_bio_ was functionally localized based on selectivity for biological motion, whereas SMG_lh_ was defined anatomically, encompassing all visually responsive voxels within the anatomical region. The left SMG was selected for analysis because it exhibited a stronger correlation with human behavioral performance than its right-hemisphere counterpart (15).

### Motion-Defined Object Stimuli

The stimuli consisted of object kinematograms, in which object shape was defined exclusively by motion cues rather than static form. Motion vectors were extracted from naturalistic videos of six object categories— three animate (human, mammal, reptile) and three inanimate (tool, ball, pendulum/swing)—following the approach described in Robert et al. (15). These motion vectors were applied to randomly positioned dot patterns, producing stimuli in which the global shape of the object was conveyed solely through coherent motion. Each category included six exemplars, resulting in a total of 36 unique kinematogram videos. This design allows the isolation of dynamic shape information from static form cues, providing a controlled way to investigate motion-defined object representations in the human visual system.

### Computational Models

We used the SlowFast architecture (18), originally trained on the Kinetics-400 dataset (22) for action recognition. The model processes visual input through two parallel pathways: a slow pathway that encodes low temporal frequency information, and a fast pathway that encodes high temporal frequency information. Occasional lateral connections allow integration of fast-varying features into the slow pathway. From this base model, we derived several variants, the intact slow pathway with cross-pathway connections (S_wx_), the fast pathway alone (F_wx_), the slow pathway with all cross-pathway connections removed (S_nox_) and S_1_, an independently trained slow-only pathway.

As a complementary model, we used DorsalNet (6), a goal-driven neural network trained on navigation tasks designed to approximate the dorsal visual pathway. Unlike SlowFast, which is optimized for categorical recognition, DorsalNet learns to extract motion features relevant for self-motion and navigation.

### Representational similarity analysis

To compare neural and model representations, we constructed representational dissimilarity matrices (RDMs) for both fMRI and model activations. For each ROI, voxel-wise responses to the 36 stimuli were extracted, averaged across repetitions and within classes, and converted into a 6 *×* 6 RDM using 1 *™* Pearson correlation. For each model and layer, activations were recorded from the ReLU outputs and converted into RDMs in the same way.

Similarity between fMRI and model RDMs was quantified using Kendall’s Tau correlation. Statistical significance was assessed using a cluster-based permutation test (23) with 1000 permutations. The test statistic was defined as the difference between the mean correlations of the observed data and a comparison condition, computed across samples. When no comparison condition was provided, correlations were tested against a zero baseline using a one-sided test; otherwise, a two-sided test was performed. Clusters of contiguous model layers were formed based on a cluster-forming threshold of *p* = 0.1 for tests. Cluster-level p-values were obtained from the permutation distribution, and layers belonging to clusters with p < 0.05 were considered statistically significant, correcting for multiple comparisons across model layers.

To account for measurement noise and estimate the upper bound of attainable correlations, we computed a noise ceiling following Nili et al. (24). The reliability of each brain region was estimated by computing Kendall’s Tau correlations between the RDM of one subject and the average RDM of all other subjects, and then averaging these reliability values across all subjects. The reliability of model RDMs was assumed to be 1, as each model layer represents a deterministic mapping from input to activation. The final noise ceiling for each region–model comparison was computed as the square root of the product of the brain and model reliabilities. This value represents the maximum correlation expected between a perfectly predictive model and the empirical neural data, given intersubject variability.

### Variance Partitioning via Partial Correlations

To determine whether networks explain the same variance of similarity or capture complementary aspects of neural representations, we computed partial correlations. Specifically, we compared the highest correlated layer of each model with ROI RDMs while controlling for variance explained by the other model. The partial correlation was computed as follows:

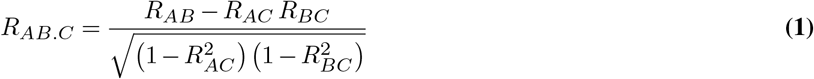

Where *R*_*AB*.*C*_ represents the unique variance in region A explained by model B beyond model C and *R*_*AB*_ denotes the Kendall’s Tau correlation between the RDMs of region A and model B. To account for the reliability of neural measurements, we computed noise ceilings using the same approach as in the RSA analysis.

### Second-Order Brain-Network similarity

To examine how brain regions and models represented object categories in a comparable space we constructed a second-order RDM (25). We first computed class-level RDMs for each brain region and model for every subject. We then compared each brain region and computational model by calculating the Kendall’s Tau correlation between the off-diagonal of their RDMs. This produced a single correlation value for every pair for each subject, capturing the similarity of their categorical representations. Across all regions and models this produced a second-order 9 *×* 9 brain-network similarity matrix for each subject that captures the pairwise similarity structure among brain regions and model layers.

To visualize this similarity structure, we applied multidimensional scaling (MDS) to each subject’s second-order similarity matrix to embed the representational relationships on a two-dimensional space using a fixed random seed. To ensure comparability across subjects, we aligned the individual MDS solutions to a common reference space using the procrustes transformation available in the Python package SciPy (26). Each subject’s region thus contributed one aligned set of coordinates, the mean position of regions across subjects on the reduced 2D space was also computed for easier comparison between models and brain regions. The resulting embeddings provided an interpretable map of representational geometries across brain regions and models. Models that aligned more closely with specific ROIs appeared spatially closer, while greater distances indicated divergent representational structures.

### Object kinematograms classification task

To test whether the models could reproduce human categorical judgments, we trained them on the same six stimulus classes used in Robert et al. (15). We replaced the final classifier of each model with a linear classification head implemented as a two-layer feedforward network: a fully connected layer (input dimension = feature size, output = 512) followed by a ReLU activation, a dropout layer (*p* = 0.5), and a final linear layer mapping to six output classes of the fMRI experiment in Robert et al. (15): human, mammal, reptile, tool, pendulum/swing, and ball. During training, only the classification head was optimized, while all other model layers were frozen. Training data were augmented by generating 10 random-dot variants of each of the 36 stimuli, producing 6 *×* 6 *×* 10 = 360 training samples. Testing was performed on the original kinematograms (the same set presented during fMRI).

We used five-fold cross-validation, where in each fold 80% of augmented stimuli were used for training and 20% held out for validation. Classification accuracy was averaged across folds to provide a robust estimate of performance.

### Behavioral Representational Analysis

Each model’s predictions on the test set were summarized in a 6 *×* 6 confusion matrix, where entry *C*_*ij*_ represents the proportion of category i stimuli classified as category j. To compare model predictions to human behavior, we converted each confusion matrix into a representational dissimilarity matrix:

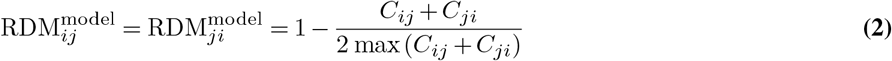

Human behavioral RDMs from Robert et al. (15) were constructed based on pairwise dissimilarity judgments. Finally, we computed Kendall’s Tau correlation between each model RDM and the behavioral RDM. This metric quantified how closely each model captured human perceptual confusions across categories.

A noise ceiling was computed to estimate the maximum attainable correlation given variability in behavioral and model data (24). Behavioral reliability was estimated using a leave-one-out (LOO) procedure, correlating each participant’s RDM with the mean of all others. Model reliability was computed analogously across cross-validation folds. The final noise ceiling was defined as the square root of the product of behavioral and model reliabilities.

## Data and Code Availability

The datasets analysed in this study were originally reported by Robert et al. (15). The computational code used to generate the results and figures is openly available at https://github.com/nastarandarjani/motion_representation.

## Competing Interests

The authors declare no competing interests.

## ACKNOWLEDGEMENTS

This work was supported by NSERC Discovery Grant (RGPIN-2023-03875) and IVADO Exploratory Project (Explo24CO-3750823649) to SB. M.V.P was supported by University of Delaware start up fund and Sloan foundation fellowship award. This research was enabled in part by support provided by (Calcul Québec) (https://www.calculquebec.ca/en/) and the Digital Research Alliance of Canada (https://alliancecan.ca/en).

## Notes

### Competing Interest Statement

The authors have declared no competing interest.

### Summary of Updates

Minor revision on text for better clarification

